# A single mutation G454A in P450 *CYP9K1* drives pyrethroid resistance in the major malaria vector *Anopheles funestus* reducing bed net efficacy

**DOI:** 10.1101/2024.04.22.590515

**Authors:** Carlos S. Djoko Tagne, Mersimine F. M. Kouamo, Magellan Tchouakui, Abdullahi Muhammad, Leon J.L. Mugenzi, Nelly M.T. Tatchou-Nebangwa, Riccado F. Thiomela, Mahamat Gadji, Murielle J. Wondji, Jack Hearn, Mbouobda H. Desire, Sulaiman S. Ibrahim, Charles S. Wondji

**Author notes:** Corresponding authors: (CSW) and (CSDT).

## Abstract

Metabolic resistance to pyrethroids is jeopardizing the effectiveness of insecticide-based interventions against malaria. The complexity of the Africa-wide spatio-temporal evolution of the molecular basis of this resistance, the major genetic drivers should be detected to improve resistance management. Here, we demonstrated that a single amino acid change G454A in the cytochrome P450 *CYP9K1* drives pyrethroid resistance in *Anopheles funestus* vector in East and Central Africa.

Polymorphism analysis revealed drastic reduction of diversity of the *CYP9K1* gene in Uganda (2014) with the selection of a predominant haplotype (90%), exhibited a G454A mutation. However, 6 years later (2020) the Ugandan 454A-*CYP9K1* haplotype was also predominant in Cameroon (84.6%), but absent in Malawi (Southern Africa) and Ghana (West Africa). *In vitro* comparative heterologous metabolism assays revealed that the mutant-type 454A-*CYP9K1* (R) allele metabolises type II pyrethroid (deltamethrin) better than the wild-type G454-*CYP9K1* (S) allele. Transgenic *Drosophila melanogaster* flies expressing the mutant-type 454A-*CYP9K1* allele were significantly more resistant to both type I and II pyrethroids than the flies expressing the wild-type G454-*CYP9K1* allele. Genotyping with a newly designed DNA-based diagnostic assay targeting the G454A replacement revealed that this mutation is strongly associated with pyrethroid resistance as mosquitoes surviving pyrethroid exposure were significantly more homozygote resistant (Odds ratio = 567, P<0.0001). Furthermore, Cone test and experimental hut trials showed that 454A-*CYP9K1* reduces the efficacy of LLINs. The resistant allele (454A) is under directional selection in Eastern and Central Africa, present but not strongly selected in Southern Africa and at very low frequency in West Africa.

This study reveals the rapid spread of P450-based metabolic pyrethroid resistance driven by *CYP9K1*, greatly reducing the efficacy of pyrethroid-based control tools. The new DNA-based assay designed here will add to the toolbox to monitor resistance in the field and improve resistance management strategies.

**Author Summary:** The complex molecular basis and genetic drivers of metabolic resistance in malaria vectors should be detected to improve resistance management. Here, we established that allelic variation by a single mutation G454A in P450 *CYP9K1* enzyme drives pyrethroid resistance in *Anopheles funestus*. Drastic reduction of diversity was noted in Ugandan female *An. funestus* samples collected in 2014, with a major haplotype (454A) already fixed but absent in other African regions. However, this Ugandan 454A-*CYP9K1* haplotype was highly selected within 6 years in *An. funestus* samples from Cameroon (Central Africa), but still absent in Ghana (West Africa) and Malawi (Southern Africa). Metabolism assays revealed that the 454A-resistant allele metabolized pyrethroid better than the susceptible G454 allele and driving higher pyrethroid resistance in transgenic *Drosophila melanogaster* flies. DNA-based diagnostics designed around the G454A-*CYP9K1* marker strongly correlates with pyrethroid resistance, reducing bed net efficacy indicating that this assay should be added to the toolbox to monitor this 454A-*CYP9K1* resistance which is rapidly spreading in *An. funestus* mosquito populations from Eastern and Central Africa.

## Introduction

Malaria is still a major public health concern despite significant progress made since the years 2000s to reduce the burden of this disease [1]. Globally, an increase in the number of cases was recently recorded from 233 million cases in 2019 to 249 million cases in 2022, with 95% of cases occurring in Africa [1]. Nevertheless, vector control mainly using Long Lasting Insecticidal Nets (LLINs) and Indoor Residual Spraying (IRS) have proven to be effective in reducing malaria burden and remains a vital component of malaria management and elimination strategy. During the past two decades, nearly 2.5 billion LLINs have been delivered to malaria endemic countries and this rapid scale-up has been by far the largest contributor to the impressive drops seen in malaria incidence and mortality since the turn of the century [2]. These control tools were attributed more than 70% of the decrease in malaria mortality and helped prevent more than 663 million clinical cases of malaria between 2000 and 2015 [3].

Unfortunately, Africa deviates from the set trajectory of the global technical strategy (GTS) milestones to eradicate malaria by 2030 [2]. This is attributed to various factors including the continuous spread and now escalation of resistance to pyrethroid insecticide by malaria vectors, which is increasingly reported in different regions across Africa [4–11], presenting greater risk of vector control failure [12]. Moreover, recent studies highlighted that pyrethroid-resistant mosquitoes present a serious challenge to the efficacy of current standard pyrethroid LLINs [8,13–16]. Studies to decipher survival strategies developed by resistant malaria vectors have reported increased expression of detoxification genes mainly cytochrome P450s, glutathione-s transferases and carboxylesterases, their overactivity and mutations [16–18], as well as insecticide target site modifications inhibiting productive insecticide binding, either in the voltage gated sodium channels, knockdown resistance (*kdr*) [19,20] or in the acetylcholine esterase receptor (Ace-1) gene [21], etc, as the major resistance mechanisms. Other mechanisms include changes in behaviour to avoid insecticide contact [22] and reduced insecticide penetration through increased production of cuticular hydrocarbon [23]. Knockdown resistance highly prevalent and driving resistance in *An. gambiae* has been absent in *An. funestus* from most African regions [24,25] and could indicate a more pronounced role of metabolic-based resistance.

Previous transcriptional profiling studies have detected key *An. funestus* cytochrome P450 genes with evidence of differential expressions that could be linked to pyrethroid resistance [15,16,26]. However, sharp regional contrast was reported with different *CYP* genes overexpressed in different African regions [16,26]. Among these genes were the duplicated *CYP6P9a/b* and *CYP6P4a/b* respectively overexpressed in *An. funestus* mosquitoes from Southern (Malawi and Mozambique) and West (Ghana) African regions, the *CYP325A* was overexpressed in central Africa (Cameroon) and the *CYP9K1* was reported to be the most up regulated gene in resistant mosquitoes from Eastern Africa (Uganda) [16,26]. The molecular bases of resistance mediated by the above P450s are gradually been deciphered revealing that *CYP325A* contributes to resistance to both type I and II pyrethroid insecticide in central Africa [27], while functional characterisation has also shown that allelic variants of *CYP6P9a* and *CYP6P9b* are major drivers of type I and type II pyrethroid resistance mainly in Southern Africa [28,29]. Further studies targeting the Promoter and intergenic regions confirmed these duplicated genes are the main drivers of pyrethroid resistance in *An. funestus* resulting to the detection of the first P450-based molecular markers [15,16,30]. However, the resistance driven by the *CYP6P9a/b* genes are limited to southern African region and the assays designed around the detected markers of these genes are mainly used to track resistance in southern Africa. Hence, there is an urgent need to detect new molecular markers driving resistance in other African regions to facilitate resistance monitoring and management in these regions.

In this regard, an Africa-wide whole genome scan and targeted enrichment with deep sequencing of permethrin-resistant *An. funestus* mosquitoes across Africa reported reduced genetic diversity with signature of directional selection and gene duplication on the X-chromosome spanning the *CYP9K1* locus [31,32]. We further analysed the genetic diversity around the *CYP9K1* gene Africa-wide and identified a glycine to alanine amino acid change on codon 454 that was fixed in Uganda, while at very low frequencies in other African regions (Cameroon and Malawi) [31]. Here, we investigated the genetic polymorphism of the *CYP9K1* gene in *An. funestus* across Africa. We then used comparative *in vitro* heterologous metabolism assay and *in vivo* transgenic *Drosophila* fly approach to assess the contribution of allelic variation and overexpression of this gene to the observed resistance. Furthermore, we designed a simple DNA-based diagnostic assay around the G454A mutation that we used to assess the impact of this marker on the efficacy of LLINs using cone test and experimental hut trials (EHTs) and also, to track the spread of this 454A-*CYP9K1* resistant marker across Africa.

## Results

### Africa-wide temporal genetic variability of *CYP9K1* gene detected a rapidly spreading 454A dominant haplotype

Two sets of samples were analysed across different African regions; Eastern Africa (Uganda mosquitoes collected in 2014 and 2020); Central Africa (Cameroon mosquitoes collected in 2014 and 2020); Southern Africa (Malawi mosquitoes collected in 2014 and 2020) and West Africa (Ghana mosquitoes collected in 2020). Genetic analysis of *CYP9K1* from 2014 samples revealed reduced diversity in Uganda 2014 samples, with 35 substitution sites and very low haplotype diversity (h_d_ = 0.19) (Table 1, Fig 1A). Contrasting pattern was observed in Cameroon 2014 samples which were highly diversified with 123 substitution sites and very high haplotype diversity (h_d_ = 0.99). Interestingly, reduced genetic diversity of *CYP9K1* was noticed in samples from Cameroon collected in 2020 exhibiting a pattern similar to Ugandan samples from 2014 and 2020. Cameroon 2020 and Uganda 2020 samples registered zero missense substitutions, both generating low haplotype and nucleotide diversities (h_d_ = 0.295, π = 0.00019 for Cameroon 2020 and h_d_ = 0.143, π = 0.00009 for Uganda 2020). However, samples from Malawi 2020 (h_d_ = 0.974, π = 0.00820) and Ghana 2020 (h_d_ = 0.873, π = 0.0046) registered higher haplotype and nucleotide diversities (Table 1, Fig 1A and B). Moreover, analysis of all the 2020 sequences yielded a total of 55 substitution sites with 9 nonsynonymous substitutions, mainly contributed by Malawi (2 substitutions), Ghana (2 substitutions) and FANG (5 substitutions).

**Fig 1.**
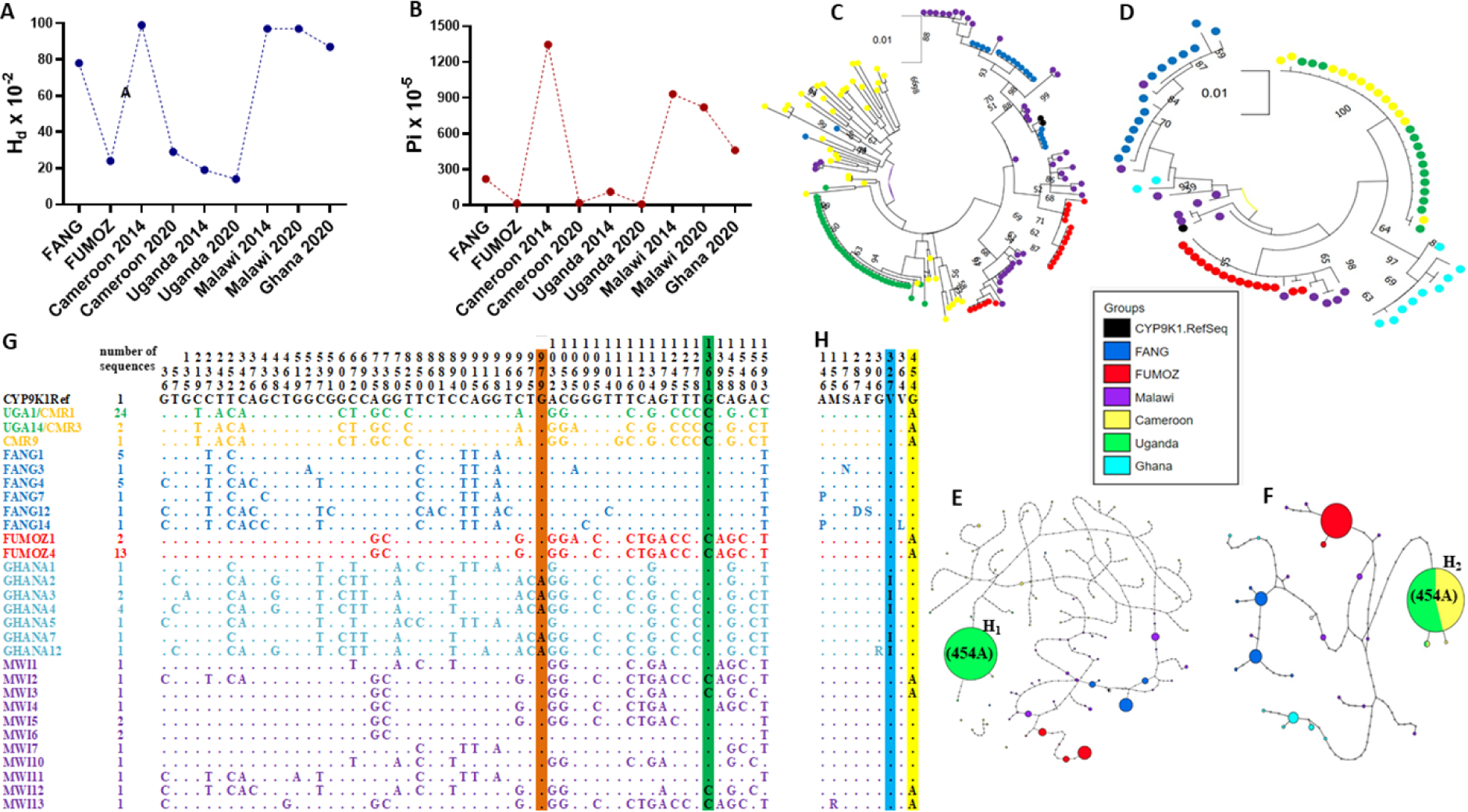
Africa-wide temporal genetic variability and haplotype representation of *CYP9K1* gene. Spatio-temporal variation of (**A**) Haplotype diversity (H_d_) and (**B**) nucleotide diversity of *CYP9K1* showing signatures of strong directional selection in Uganda and Cameroon between 2014 and 2020. (**C**) Maximum likelihood tree of *CYP9K1* coding length across Africa using 2014 samples. (**D**) Maximum likelihood tree of *CYP9K1* coding length across Africa using 2020 samples. (**E**) Haplotype network representation of *CYP9K1* coding across Africa using 2014 samples showing dominant haplotype shared between Uganda samples. (**F**) Haplotype network representation of *CYP9K1* coding across Africa using 2020 samples showing dominant haplotype shared between Cameroon 2020 and Uganda 2020. (**G**) polymorphic positions of *CYP9K1* cDNA sequences in 2020 samples showing nucleotide changes, the guanine to adenine nucleotide at position 979 (highlighted in orange colour) and guanine to cytosine nucleotide at position 1361 (highlighted in green colour). (**H**) polymorphic positions of *CYP9K1* amino acid sequences 2020 samples highlighting the V327I (highlighted in blue) and G454A (highlighted in yellow) amino acid changes.

**Table 1:**
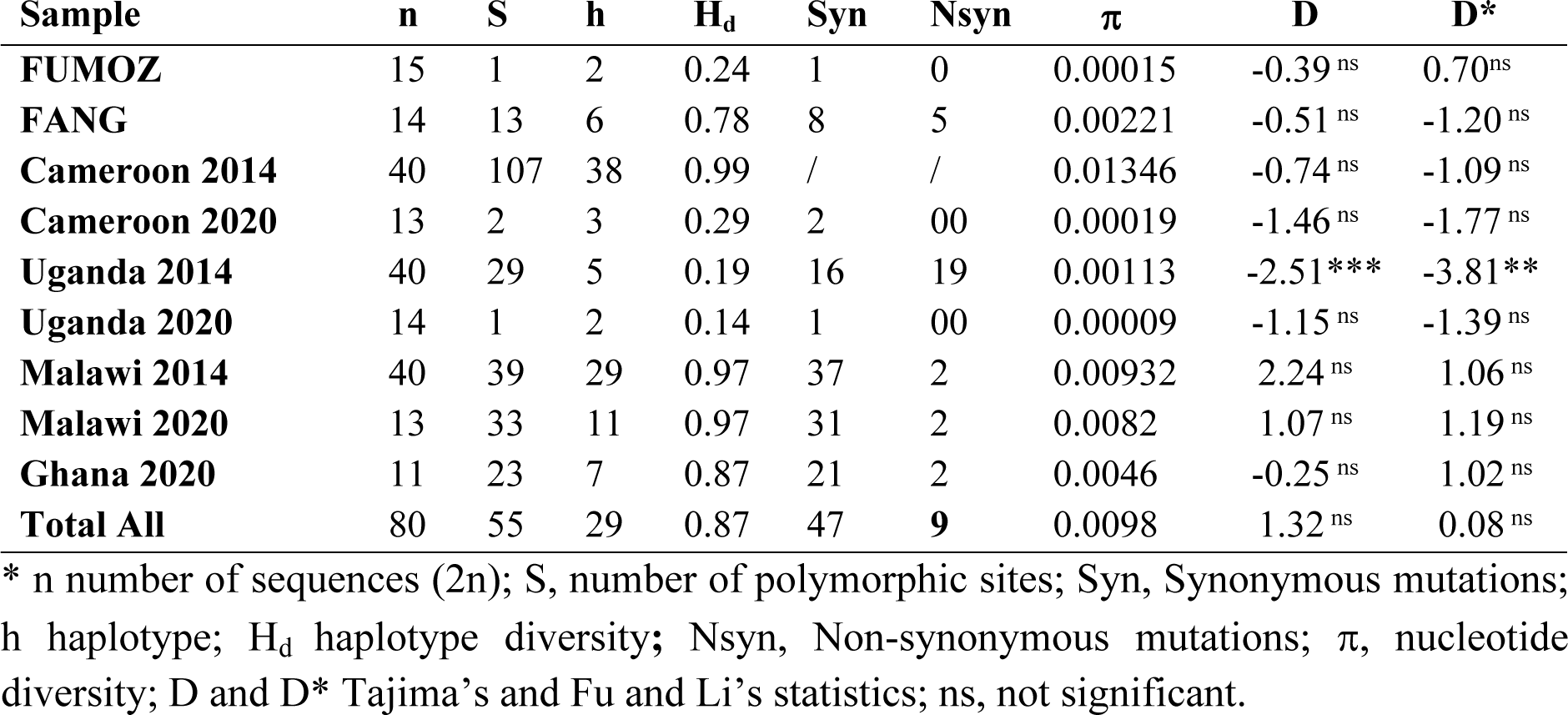
Africa-wide Polymorphism parameters of CYP9K1.

| Sample | n | S | h | H<sub>d</sub> | Syn | Nsyn | π | D | D* |
| --- | --- | --- | --- | --- | --- | --- | --- | --- | --- |
| FUMOZ | 15 | 1 | 2 | 0.24 | 1 | 0 | 0.00015 | -0.39<sup>ns</sup> | 0.70<sup>ns</sup> |
| FANG | 14 | 13 | 6 | 0.78 | 8 | 5 | 0.00221 | -0.51<sup>ns</sup> | -1.20<sup>ns</sup> |
| Cameroon 2014 | 40 | 107 | 38 | 0.99 | / | / | 0.01346 | -0.74<sup>ns</sup> | -1.09<sup>ns</sup> |
| Cameroon 2020 | 13 | 2 | 3 | 0.29 | 2 | 00 | 0.00019 | -1.46<sup>ns</sup> | -1.77<sup>ns</sup> |
| Uganda 2014 | 40 | 29 | 5 | 0.19 | 16 | 19 | 0.00113 | -2.51*** | -3.81** |
| Uganda 2020 | 14 | 1 | 2 | 0.14 | 1 | 00 | 0.00009 | -1.15<sup>ns</sup> | -1.39<sup>ns</sup> |
| Malawi 2014 | 40 | 39 | 29 | 0.97 | 37 | 2 | 0.00932 | 2.24<sup>ns</sup> | 1.06<sup>ns</sup> |
| Malawi 2020 | 13 | 33 | 11 | 0.97 | 31 | 2 | 0.0082 | 1.07<sup>ns</sup> | 1.19<sup>ns</sup> |
| Ghana 2020 | 11 | 23 | 7 | 0.87 | 21 | 2 | 0.0046 | -0.25<sup>ns</sup> | 1.02<sup>ns</sup> |
| Total All | 80 | 55 | 29 | 0.87 | 47 | 9 | 0.0098 | 1.32<sup>ns</sup> | 0.08<sup>ns</sup> |

\* n number of sequences (2n); S, number of polymorphic sites; Syn, Synonymous mutations; h haplotype; H<sub>d</sub> haplotype diversity; Nsyn, Non-synonymous mutations; π, nucleotide diversity; D and D\* Tajima's and Fu and Li's statistics; ns, not significant.

Interestingly, Ugandan 2014 samples clustered together forming a dominant clade, not observed in Cameroon 2014 and Malawi 2014 with higher diversities and no signs of directional selection (Fig 1C). However, analysis of 2020 samples revealed a clustering between Cameroon 2020 and Uganda forming the same dominant clade, different from other African regions, Southern Africa (Malawi 2020) and West Africa (Ghana 2020) that clustered independently, forming minor clades per region (Fig 1D).

Furthermore, haplotype network analysis of *CYP9K1* in 2014 samples revealed a dominant haplotype (H_1_) present at 90% frequency in Uganda 2014 samples with a total of 5 haplotypes, not observed in other African regions where the haplotype was completely absent (0%), like in Cameroon 2014 and Malawi 2014 registering total of 38 and 29 haplotypes respectively (S1 Fig). Cameroon 2014 samples were highly diverse forming singleton haplotypes with no dominant haplotype detected (Fig 1E). Haplotype networking using 2020 samples revealed Cameroon 2020 and Uganda 2020 samples both shared the same dominant haplotype (H_2_) (Fig 1F). This was the same haplotype found at very high frequency in Uganda 2014 and completely absent in Cameroon 2014 samples. By 2020 the H_2_ haplotype has become predominant in Cameroon with a frequency nearing fixation at 84.6%, while absent in Southern Africa (Malawi) and West Africa (Ghana). Furthermore, haplotype network analysis of 2020 samples generated a total of 29 haplotypes mainly distributed between more diverse samples from Malawi (11 haplotypes), Ghana (7 haplotypes) and laboratory susceptible strain FANG (6 haplotypes) (Fig 1E, Table 1). Fewer haplotypes were detected in samples from Uganda (2 haplotypes) and Cameroon (3 haplotypes), while strong allelic variations were observed between samples from Southern (Malawi) and West (Ghana) African regions, each forming distinct minor haplotypes.

Sequence examination identified a point mutation guanine (G) to cytosine (C) nucleotides at position 1361 (Fig 1G), leading to replacement of glycine (G) to Alanine (A) on codon 454 (Fig 1H). This mutation was fixed in Uganda in 2014 (40/40), at low frequency in Cameroon in 2014 (9/40) and Malawi in 2014 (13/40) (Fig S1). However, analysis of 2020 sequences revealed the mutation was fixed in Uganda (14/14, 100%) and Cameroon (13/13, 100%) cDNA samples, but still at low frequency in Malawi (4/13, 30.7%) and completely absent in both Ghana (0/11, 0%) and FANG (0/14, 0%) (Fig 1H). Another point mutation guanine to adenine (A) nucleotides at position 979, resulting to amino acid change valine (V) to isoleucine (I) on codon 327 detected at high frequency (9/11, 81.8%) in samples from Ghana (West Africa) and absent in samples from Cameroon, Malawi and Uganda (Fig 1H).

### Comparative *in vitro* assessment of metabolic activity of *CYP9K1* allelic variants

#### Expression pattern of recombinant *CYP9K1* alleles

Co-expression of recombinant *CYP9K1* with *CPR* revealed optimal expression between 22-24 h post induction with 1 mM IPTG and 0.5 mM δ-ALA. Importantly, both alleles of *CYP9K1* when complexed with standard P450 carbon monoxide (CO) generated similar difference spectra with absorbance peaks at 450nm wavelength (Fig 2A and B), with comparable concentrations of 3.27 μM for mutant allele (454A-*CYP9K1*) and slightly higher concentration of 3.67 μM for the wild allele (G454-*CYP9K1*).

**Fig 2.**
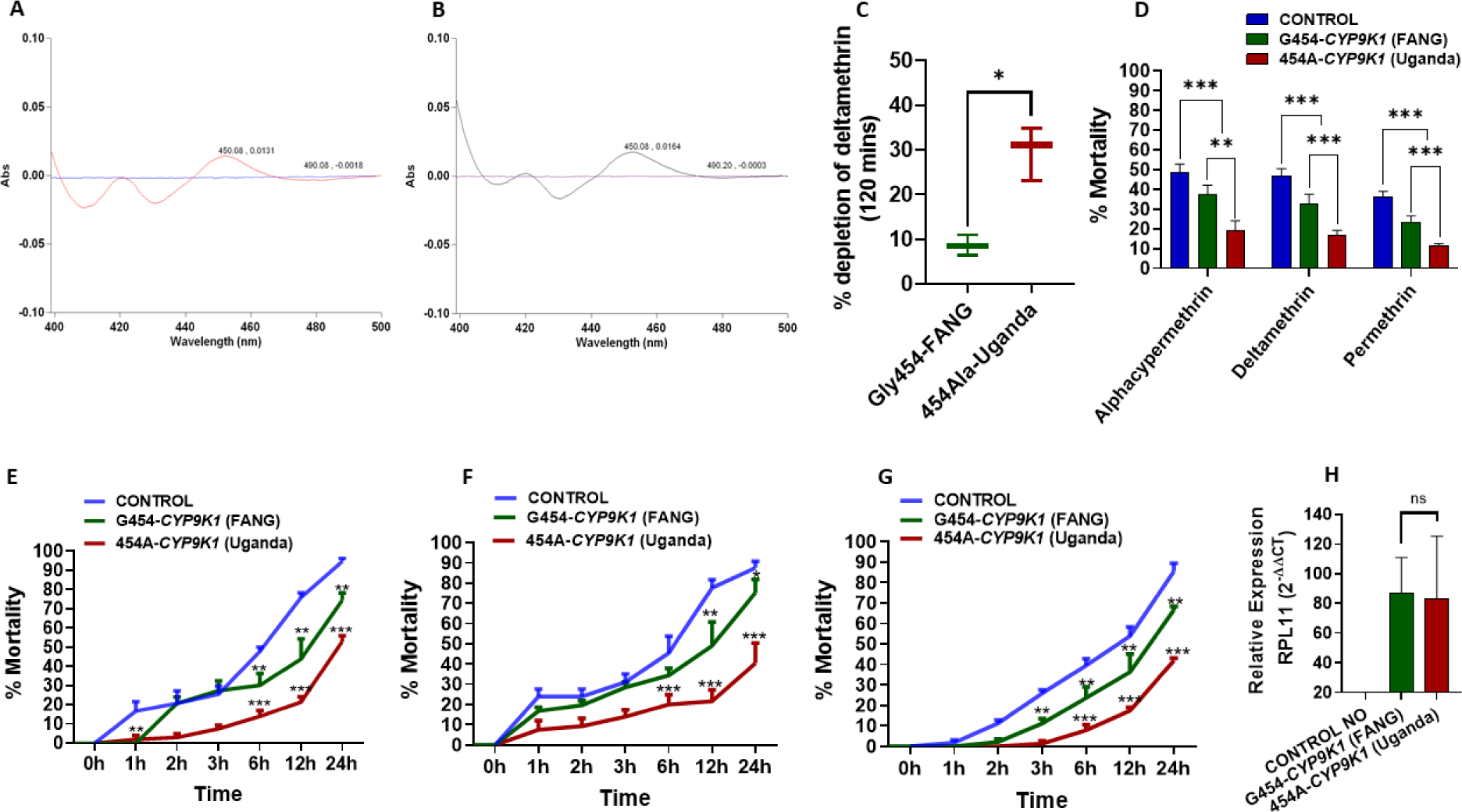
Comparative *in vitro* and *in vivo* assessment of *CYP9K1* alleles to confer pyrethroid resistance. (**A**) Co-difference spectra of membranes expressing recombinant *CYP9K1* mutant-type (454A) allele from Uganda and Cameroon and (**B**) Wild-type (G454) allele from Laboratory susceptible strain FANG. (**C**) Percentage depletion of 20 µM deltamethrin by recombinant *CYP9K1* membranes. Horizontal bar represents the mean percentage depletion, and the error bars represent the 95% confidence intervals of the mean. The p value is calculated using a Welch’s t test. Results are average of three replicates compared between the two groups. (**D**) Average percentage mortality of *CYP9K1* allelic variants 24 hours after exposure to pyrethroid insecticide; Comparative mortalities of F_1_ transgenic progenies of crosses between Actin5C-GAL4 and *UAS-CYP9K1* on exposure to (**E**) 0.0007% alphacypermethrin; (**F**) 0.2% deltamethrin and (**G**) 4% permethrin; (**H**) Comparative relative expression of G454-*CYP9K1* and 454A-*CYP9K1* in *Drosophila* flies. Data are represented as mean ± SEM: *P<0.05, ** P<0.01 and *** P<0.001.

Preliminary metabolism assays revealed that the recombinant 454A-*CYP9K1* depleted higher amount (29.7%) of deltamethrin compared to the G454-*CYP9K1* allele that depleted only 8.5% of this type II pyrethroid (Fig 2C). Depletion of deltamethrin was 21.10% significantly greater (SE = 3.68) in the mutant-type allele than in the wild-type allele (*t*-test: *t* = 5.73; df = 2.59; P = 0.01) demonstrating that mutant-type 454A-*CYP9K1/CPR* allele is better metabolizer of deltamethrin *in vitro* than the wild-type allele G454-*CYP9K1/CPR*. However, as previously established in previous studies both alleles did not deplete permethrin, suggesting lack of metabolic activity towards this type I pyrethroid.

### Comparative assessment of *CYP9K1* alleles to confer pyrethroid resistance using GAL4/UAS transgenic expression in *D. melanogaster*

#### Contact bioassays with pyrethroid insecticides

Bioassays with alphacypermethrin, deltamethrin and permethrin revealed that transgenic *Drosophila melanogaster* flies overexpressing both alleles confer pyrethroid resistance, with the flies expressing the mutant 454A-*CYP9K1* allele significantly more resistant to all pyrethroids tested than those expressing G454-*CYP9K1* wild allele and control. The type 454A-*CYP9K1* flies exhibited significantly lower mortalities on average, of 18.26%±5.33 (P<0.0004), 16.83%±2.41 (P<0.0006) and 11.44%±1.15 (P<0.0006) respectively, for alphacypermethrin, deltamethrin and permethrin compared to G454-*CYP9K1* flies, with mortalities for the above pyrethroids respectively of 37.27%±4.73, 32.77%±4.76 and 23.36%±3.24 (Fig 2D). Compared to control groups with average mortalities of 48.26%±4.49 (P<0.0055), 46.87%±3.66 (P = 0.0009), and 36.36%±2.69 (P<0.0001), respectively for alphacypermethrin, deltamethrin and permethrin.

##### Bioassays with alphacypermethrin

Time-course assay revealed that the transgenic 454A-*CYP9K1* were more resistant to alphacypermethrin, with mortalities of less than 10% (P<0.0001), 20% (P = 0.002) and 30% (P = 0.0066) in the first 2h, 6h and 12h, respectively, compared with higher mortalities at the same time point recorded in the G454-*CYP9K1* flies (20%, 30% and 45% mortalities, respectively), while both experimental flies exhibited significantly lower mortalities when compared with control flies (Fig 2E).

##### Bioassays with deltamethrin

Similar to alphacypermethrin, bioassay results with deltamethrin revealed lower mortality in flies expressing mutant allele (454A) than those expressing wild allele (G454) and control flies. Transgenic 454A-*CYP9K1* flies registered mortalities around 10% (P = 0.0057), 20% (P = 0.0038), less than 30% (P = 0.0054) in the first 2hrs, 6hrs and 12hrs respectively, compared to significantly higher mortalities recorded in flies expressing the wild allele (with 20%, 35%, 50% mortalities respectively at the same time). Interestingly, flies expressing either the mutant (454A) or wild allele (G454) registered significantly lower mortalities compared to control flies (Fig 2F).

##### Bioassays with permethrin

Bioassays with permethrin resulted to similar patterns observed with type II pyrethroids, with lower percentage mortality rates for flies expressing the *CYP9K1* mutant allele when compared to the mortality rate in flies expressing the wild allele. Transgenic 454A*-CYP9K1* flies registered less than 2% (P = 0.0009) mortality in the first 3h after exposure to permethrin, less than 10% (P = 0.0049), less than 20% (P = 0.0219), and 40% (P<0.0001) respectively after 6hrs, 12hrs and 24hrs compared to flies expressing the G454*-CYP9K1* wild allele that registered higher mortalities (around 22%, 32%, and 62% mortalities respectively at the same time) (Fig 2G).

### Validation of overexpression of transgenes by qRT-PCR

Overexpression of *CYP9K1* was confirmed in the F_1_ progenies of the GAL4/UAS crosses using RT-qPCR. Both alleles were overexpressed in the F_1_ progenies when compared to the control (generated from the crossing between the UAS line without the gene and the Act5C-GAL4 driver line) (Fig 2H). Interestingly, no significant difference (t-test = 0.14, df = 3.15) was observed in the expression levels between the G454-*CYP9K1* flies (FC = 87.3) and the 454A-*CYP9K1* flies (FC = 83.17).

### Design of G454A-*CYP9K1* diagnostic assay and correlation with pyrethroid resistance

To detect and track resistance driven by 454A-*CYP9K1* allele, two PCR-based assays: an allele specific PCR (AS-PCR) and a probe-based locked nucleic acid (LNA) were successfully designed (Fig 4A and B), targeting the glycine (1361-G**G**A) to alanine (1361-G**C**A) codon 454 mutation. Using genomic DNA samples extracted from Uganda female *An. funestus* mosquitoes (Mayuge F_0_ 2021), these assays revealed that the 454A mutation was present at fixation with 100% frequency, with all the samples harbouring the resistant allele exhibiting a single band at 216 bp corresponding to homozygote resistant genotype 454A/A-*CYP9K1* (RR) and a common band around 639 bp on gel image. This was confirmed with LNA-assay where all samples clustered together on the y-axis in the LNA dual scatter plot. On the other hand, all the laboratory susceptible samples FANG genotyped were homozygote wild-type G/G454-*CYP9K1* genotype (SS), with the wild-type band size around 434bp with a common band around 639bp on gel image and clustering together on the x-axis in the LNA dual scatter plot (S2A and B Fig).

To obtain a more robust genotype segregation FANG strain were crossed with Mayuge mosquitoes and reared to F_5_ generation, which were genotyped with Figure 4A and B depicting better segregation of genotypes (Fig 3A and B).

**Fig 3.**
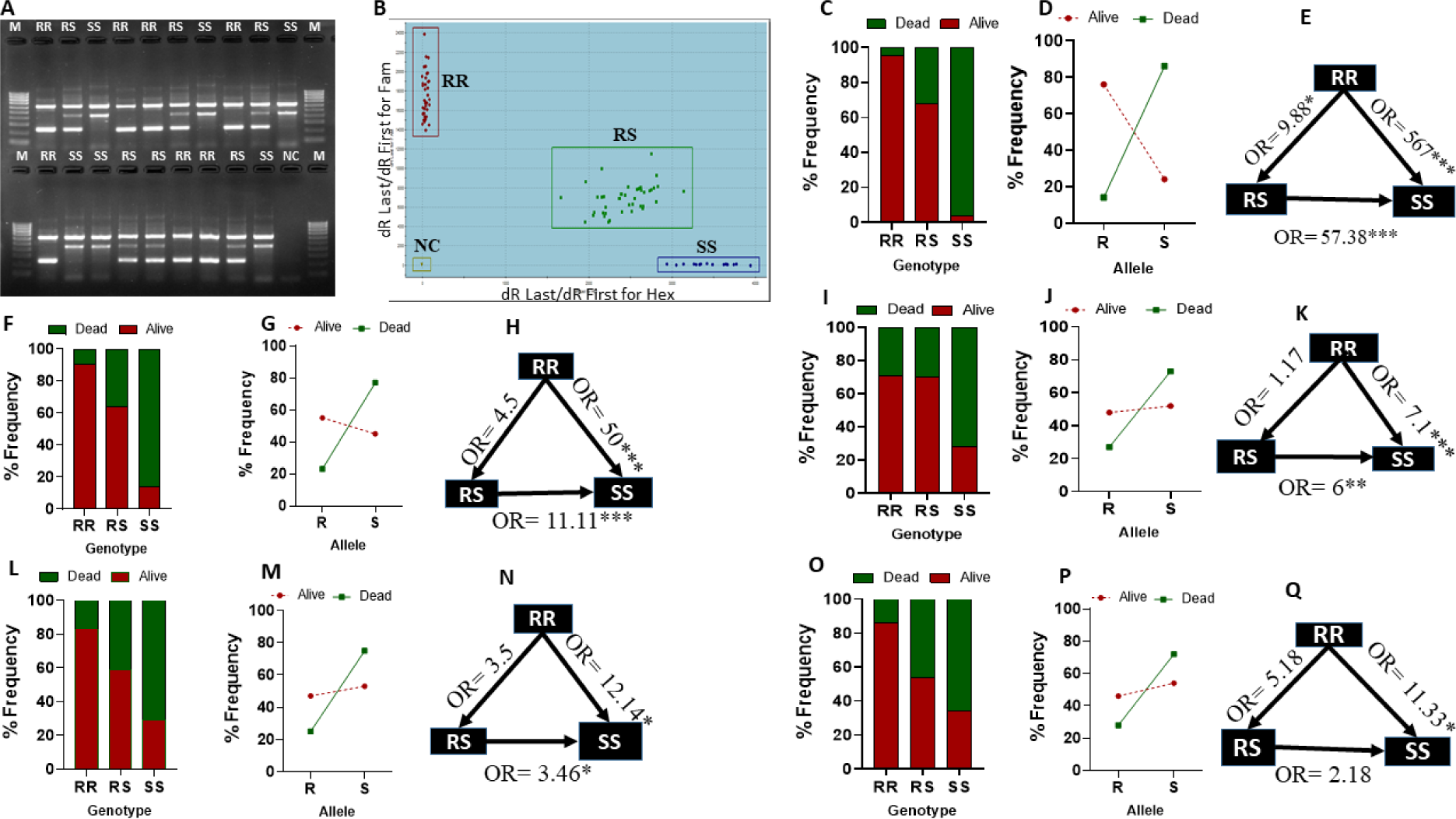
Design of DNA-based diagnostic assay for G454A-*CYP9K1* genotyping and correlation with pyrethroid resistance phenotype. (**A**) Agarose gel image of G454A-*CYP9K1* AS-PCR showing homozygote mutant-type 454A/A-*CYP9K1* (RR, with band size 216 bp and a common band 639 bp), heterozygote G/A454-*CYP9K1* (RS, band sizes 216 bp and 434 bp with at common band 639 bp), homozygote wild-type G/G454-*CYP9K1* (SS, with band size 434 bp plus a common band 639 bp) genotypes, (**B**) Dual colour scatter plot of *CYP9K1*-probe based locked-nucleic acid (LNA) showing homozygote mutant-type 454A/A-*CYP9K1* (RR) genotypes clustered on the y-axis and homozygote wild-type G/G454-*CYP9K1* (SS) genotypes clustered on the x-axis, heterozygotes (RS) in between the x-axis and y-axis and negative controls (NC), **(C)** G454A-*CYP9K1* percentage genotype frequency distribution in FANG x Mayuge F_5_ permethrin alive and dead female mosquitoes. **(D)** Percentage allele frequency distribution FANG x Mayuge F_5_ permethrin alive and dead female mosquitoes. **(E)** Estimation of odds ratio (OR) and associated significance between different *CYP9K1* genotype in permethrin alive and dead FANG x Mayuge F_5_ female mosquitoes. The arrow within the triangle indicates the direction of OR estimation and ORs are given with asterisks indicating level of significance. **(F)** G454A-*CYP9K1* genotype frequency distribution in FANG x Mibellon F_3_ alphacypermethrin alive and dead female mosquitoes. **(G)** Percentage allele frequency distribution FANG x Mibellon F_3_ alphacypermethrin alive and dead female mosquitoes. **(H)** Estimation of odds ratio (OR) and associated significance between different *CYP9K1* genotype alphacypermethrin alive and dead female mosquitoes. **(I)** G454A-*CYP9K1* genotype frequency distribution in Elende 2022 F_1_ alphacypermethrin alive and dead female mosquitoes. **(J)** Allelic frequency distribution in Elende 2022 F_1_ alive and dead to alphacypermethrin exposure. **(K)** Estimation of odds ratio (OR) and associated significance between different *CYP9K1* genotype in Elende 2022 F_1_ alphacypermethrin alive and Dead. **(L), (M),** (**N**) are respectively the same as **(I)**, **(J)**, **(K)** using Elende 2022 F_1_ permethrin alive and dead and **(O)**, **(P)**, **(Q)** are respectively the same as **(L)**, **(M)**, **(N)** using Elende 2022 F_1_ deltamethrin alive and dead.

**Fig 4.**
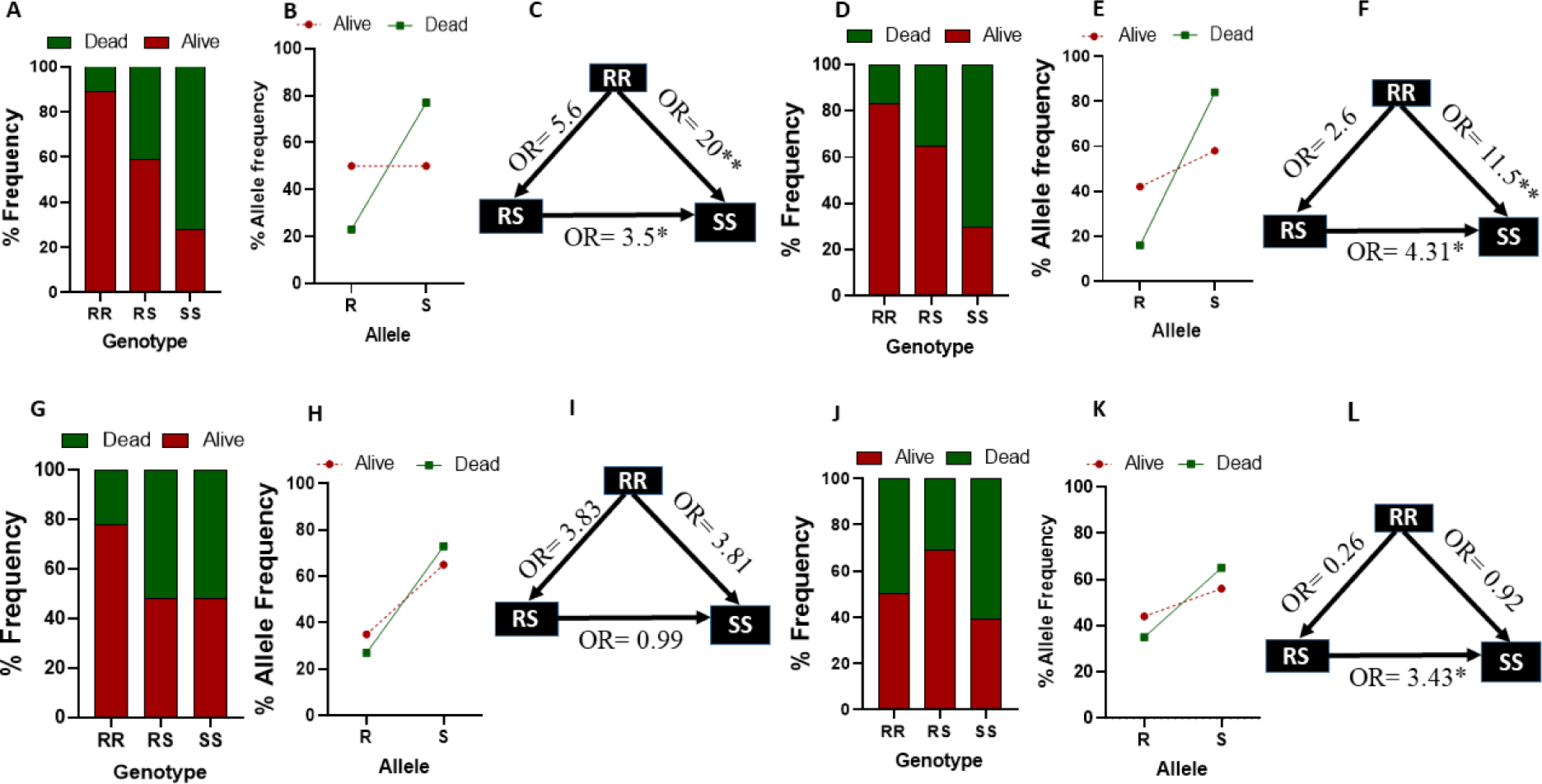
Correlation between the G454A-*CYP9K1* mutation and resistance to LLINs using cone test and EHTs samples. **(A)** *CYP9K1* percentage Genotype frequency distributions in Elende F_1_ alive and dead to Royal sentry using cone test. **(B)** *CYP9K1* percentage allele frequency distributions in Elende F_1_ alive and dead to Royal sentry using cone test. **(C)** Estimation of odds ratio (OR) and associated significance between different *CYP9K1* genotypes and the ability to survive exposure to Royal sentry using cone test. The arrow within the triangle indicates the direction of OR estimation and ORs are given with asterisks indicating level of significance. **(D), (E)**, **(F)** are respectively the same as **(A)**, **(B)**, **(C)** using Elende 2022 F_1_ alive and Dead to Olyset exposure. **(G)** *CYP9K1* percentage Genotype frequency distributions in Elende 2022 F_0_ free-flying alive and dead to Royal sentry using EHT. **(H)** *CYP9K1* percentage allele frequency distributions in Elende 2022 F0 free-flying alive and dead to Royal sentry using EHT. **(I)** Estimation of odds ratio (OR) and associated significance between different *CYP9K1* genotypes and the ability to survive exposure to Royal sentry using EHT. The arrow within the triangle indicates the direction of OR estimation and ORs are given with asterisks indicating level of significance. **(J)**, **(K)**, **(L)** are respectively the same as **(G)**, **(H)**, **(I)** using Elende 2022 F_0_ free-flying alive and Dead to PermaNet 3.0 exposure using EHTs.

### Assessment of association between the G454A-*CYP9K1* mutation and pyrethroid resistance

#### The G454A-*CYP9K1* strongly correlates with pyrethroid resistance phenotype

To assess correlation between the G454A mutation and pyrethroid resistance phenotype, a WHO tubes bioassays was conducted with 0.75% permethrin and using F_5_ hybrid progenies generated from crosses between FANG and Mayuge *An. funestus*. Female mosquito samples highly resistant (78%±4 mortalities, alive following 1 h permethrin 0.75% exposure) and highly susceptible (26%±2 mortalities, dead following 30 minutes exposure) were genotyped. A clear distribution of genotype was observed with 55% of permethrin alive female mosquitoes being homozygote resistant (RR = 22/40), 42.5% heterozygote (RS = 17/40), and only a single alive female was homozygote susceptible (SS = 1/40) genotype. In Contrary, 72.5% of the permethrin-dead female mosquitoes (SS = 29/40) harboured the homozygote susceptible genotype, 25% were heterozygotes (10/40), with only a single dead female homozygote resistant (1/40) (Fig 3C).

Allele frequency distribution revealed a strong positive association between mosquitoes harbouring the mutant allele (R) and ability to survive permethrin exposure (OR: 19.45; CI: 9.25-38.48; P<0.0001) (Fig 3D). Mosquitoes carrying the homozygote resistant genotype (RR) significantly survived exposure to permethrin than heterozygotes (RS) (OR: 9.88; CI: 1.485-114.1; P = 0.025, Fisher’s exact test) (S1 Table) and the homozygote susceptible (SS) (OR: 567; CI: 34.03-5831; P<0.0001, Fisher’s exact test) genotype (Fig 3E). Furthermore, the heterozygotes (RS) were predominant in the alive mosquitoes (68%) than in the dead (32%) and revealing a strong association between harbouring the heterozygote genotype (RS) and ability to survive than the homozygote susceptible genotype (SS) (OR: 57.38; CI: 7.47-617.6; P<0.001, Fisher’s exact test).

A similar pattern was observed for female *An. funestus* mosquitoes exposed to alpha-cypermethrin alive after 1 h exposure (highly resistant) and dead after 10 minutes exposure (highly susceptible), with 90% of the homozygote mutant genotype (RR) recorded for the alive female mosquitoes versus 10% frequency in the dead (Fig 3F). Moreover, female mosquitoes alive to alphacypermethrin were scored with 55% of the mutant (R) allele, while 77% of the wild (S) allele was recorded in the dead female *An. funestus* mosquitoes (Fig 3G). Statistical analysis revealed an increase survivorship to alphacypermethrin in mosquitoes harbouring homozygote mutant-type genotype (RR) when compared to the homozygote wild-type genotype (SS) (OR: 50; CI: 5.23-566.4; P<0.0001, Fisher’s exact test) (S1 Table). Moreover, significant increase in survival was registered for the heterozygote (RS) genotype when compared to the homozygote wild genotype counterpart (OR: 11.11; CI: 3.31-32.28; P<0.0001, Fisher’s exact test) (Fig 3H).

#### 454A-CYP9K1 mutant-type allele (S) correlates with resistance using field samples

We further assessed the association between the G454A-*CYP9K1* marker and resistance to pyrethroid using field samples from Elende (F_1_) collected in 2022 (Central Cameroon). WHO tube test using Elende F_1_ samples with 0.75% permethrin revealed 59.55±1.7% mortality after 60 minutes exposure, 0.05% deltamethrin and 0.05% alphacypermethrin revealed 70.21±2.5% and 34.4±2.8% mortalities respectively after 60 minutes exposure.

Genotype frequency distribution of G454A-*CYP9K1* mutation in field samples exposed to alphacypermethrin alive and dead after 60 minutes revealed 71.4% frequency of homozygote mutant-type genotype (RR) in alive female *An. funestus* samples (5/7) and only 28.6% in dead (2/7). In the contrary, homozygote wild genotype (SS) was more abundant with 72%frequency in dead samples (18/25) than in alive (7/25) (Fig 3I). Allele frequency revealed 71.4% of the wild (S) allele in dead samples and only 27% of the mutant (R) allele in dead (Fig 3J). Statistically, a significant increase in survival ability was recorded between the homozygote mutant-type genotype (RR) compared to the homozygote wild (SS) genotype (OR: 7.1; CI: 2.2-22.64; P=0.0006, Fisher’s exact test) (S1 Table). Similarly, heterozygotes (RS) revealed significantly higher survivorship to alphacypermethrin when compared to the homozygote wild-type (SS) genotype (OR: 6; CI: 1.897-17.42; P<0.001, Fisher’s exact test) (Fig 3K).

Similar pattern was obtained for female *An. funestus* mosquito samples alive and dead to 0.75% permethrin with 83% of the homozygote mutant (RR) genotype detected in alive mosquitoes, while dead mosquitoes were more homozygote wild (SS) genotype with 71% frequency (Fig 3L). The wild (S) allele was more associated with susceptibility to permethrin, with 75% of this allele registered in dead samples (Fig 3M). Indeed, field mosquitoes bearing the homozygote mutant genotype (RR) were significantly more resistant to permethrin compared those harbouring the wild-type (SS) genotype (OR: 12.14; CI: 1.55-49.4; P<0.025, Fisher’s exact test) (Fig 3N).

Genotyping samples alive and dead to 0.05% deltamethrin revealed 86% of the homozygote resistant (RR) genotype in alive samples. In the contrary, 66% of the homozygote wild (SS) genotype scored in dead samples (Fig 3O). Percentage allele frequency distribution revealed 72% of the susceptible (SS) in the dead mosquitoes (Fig 3P). A significant increase in survival capacity to deltamethrin was recorded in mosquitoes bearing the homozygote resistant (RR) genotype compared to mosquitoes carrying the homozygote susceptible (SS) genotype (OR: 11.33; CI: 1.25-134.1; P=0.03, Fisher’s exact test) (Fig 3Q).

#### Correlation between the G454A-*CYP9K1* mutation and resistance to LLINs using WHO cone bioassays

Cone assay results with F_1_ Elende samples revealed very low mortality rate with pyrethroid only nets with mortality 8.3±2.3% for PermaNet 3.0 side (without PBO), PermaNet 2.0 registered mortality 10.53±1.5%, Royal sentry, Olyset and DuraNet registered mortalities 13.56±3.1%, 18.33±2.2% and 10.23±2.7% respectively. Nevertheless, PermaNet 3.0 top and Olyset plus, both containing PBO were the most effective nets both registering 100% mortality.

Assessment of the genotype distribution in samples alive and dead to Royal sentry net exposure revealed 89% of the homozygote mutant genotype (RR) in alive samples and oppositely, 72% of the wild genotype (SS) was registered in dead female *An. funestus* mosquito samples (Fig 4A). Moreover, 77% of the wild (S) allele was scored in dead samples (Fig 4B). A statistically significant increase in the ability to survive Royal sentry exposure was observed for mosquito samples harbouring homozygote mutant genotype (RR) compared to the homozygote wild genotype (SS) (OR: 20; CI: 2.57-230.3; P<0.0023, Fisher’s exact test) (S1 Table). Also, harbouring a single copy of the *CYP9K1* confers an increase survivorship as heterozygote mosquitoes (RS) were more in alive compared to homozygote susceptible (SS) (OR: 3.5; CI: 1.18-9.51; P<0.02, Fisher’s exact test) (Fig 4C).

Similar pattern was reported for samples alive and dead to Olyset net exposure, with 83% of the mutant genotype (RR) in alive samples and in the contrary, 70% of the wild-type (SS) genotype was registered in dead samples (Fig 4D). Furthermore, 84% of the wild-type G454-*CYP9K1* (S) allele was scored in dead samples (Fig 4E). A significant increase in the ability to survive Olyset exposure for samples carrying the homozygote mutant genotype (RR) compared to the homozygote wild-type genotype (SS) (OR: 11.5; CI: 1.48-140.0; P<0.023, Fisher’s exact test) (S2 Table). Heterozygotes (RS) were also significantly more alive than dead post exposure to Olyset when compared to homozygote wild genotype (SS) samples (OR: 4.31; CI: 1.43-12.79; P<0.014, Fisher’s exact test) (Fig 4F).

#### Correlation between the G454A-*CYP9K1* and insecticide treated bed net using EHTs

Genotype distribution of samples alive and dead to Royal sentry (both room and veranda) revealed higher frequency of the homozygote mutant (RR) genotype in alive samples (78%) and only 22% frequency in dead samples (Fig 4G). However, even though the mutant-type 454A-*CYP9K1* (R) allele frequency was higher in the alive samples (35%) than in the dead samples (27%), the wild type G454-*CYP9K1* (S) allele was more prevalent in both phenotypes with a higher frequency in dead samples (73%) than alive samples (65%) (Fig 4H). Nevertheless, statistical analysis revealed mosquitoes bearing homozygote mutant-type genotype (RR) to be 3.8 times more likely to survive exposure than homozygote wild genotype (SS) (OR: 3.81; CI: 0.82-19.29; P=0.14, Fisher’s exact test) (Fig 4I).

Genotyping female *An. funestus* samples alive and dead from huts treated with PBO-net PermaNet 3.0 revealed equal distribution of the homozygote mutant genotype (RR) in alive (50%) and dead (50%) samples. However, the heterozygote genotype (RS) was more prevalent in alive (69%) than in dead samples (31%) (Fig 4J). Similarly, the homozygote wild genotype (SS) was more represented in dead samples (61%) than in alive samples (39%) (Fig 4J). Furthermore, allele frequency distribution revealed higher frequency of mutant (R) allele in alive samples (44%) than dead samples (35%), while the wild-type allele was more prevalent in dead samples (65%) than alive samples (56%) (Fig 4K). Interestingly, heterozygotes (RS) registered a significant increase survivorship to PermaNet 3.0 compared to the homozygote wild genotype (SS) (OR: 3.43; CI: 1.5-7.64; P=0.0057, Fisher’s exact test) (Fig 4L, S2 Table).

### Africa-wide spatio-temporal distribution of the G454A-*CYP9K1* marker

Application of this novel diagnostic assay to assess the distribution and temporal evolution of G454A-*CYP9K1* marker in female *An. funestus* samples collected at different time points in different regions in Africa revealed marked differences according to regions (Fig 5A). Investigation of the percentage genotype of G454A-*CYP9K1* marker in samples from West Africa revealed 83% and 97% of the samples were homozygote wild genotype (SS) in Ghana 2021 and Benin 2021 respectively, with no female *An. funestus* mosquito being homozygote mutant genotype (RR) (Fig 5A). However, the marker was detected at moderate frequencies for the homozygote mutant genotype (RR) in samples from southern Africa, 28% frequency in Zambia 2014, 10% in Mozambique 2020 and 27% in Malawi 2021. Genotyping samples from Eastern Africa revealed the homozygote mutant genotype (RR) was present at 58% in Tanzania in female *An. funestus* mosquito samples collected in 2018. The marker is present at fixation, 100% frequency in Uganda 2021 with all female *An. funestus* mosquito samples genotyped bearing the homozygote mutant genotype (RR). This was similar to samples from Central Africa, Cameroon (Mibellon 2021) where the homozygote mutant genotype (RR) was detected near fixation at 90% frequency (Fig 5A). Nevertheless, the homozygote mutant genotype (RR) was detected at lower frequency of 30% in DRC 2021.

**Fig 5.**
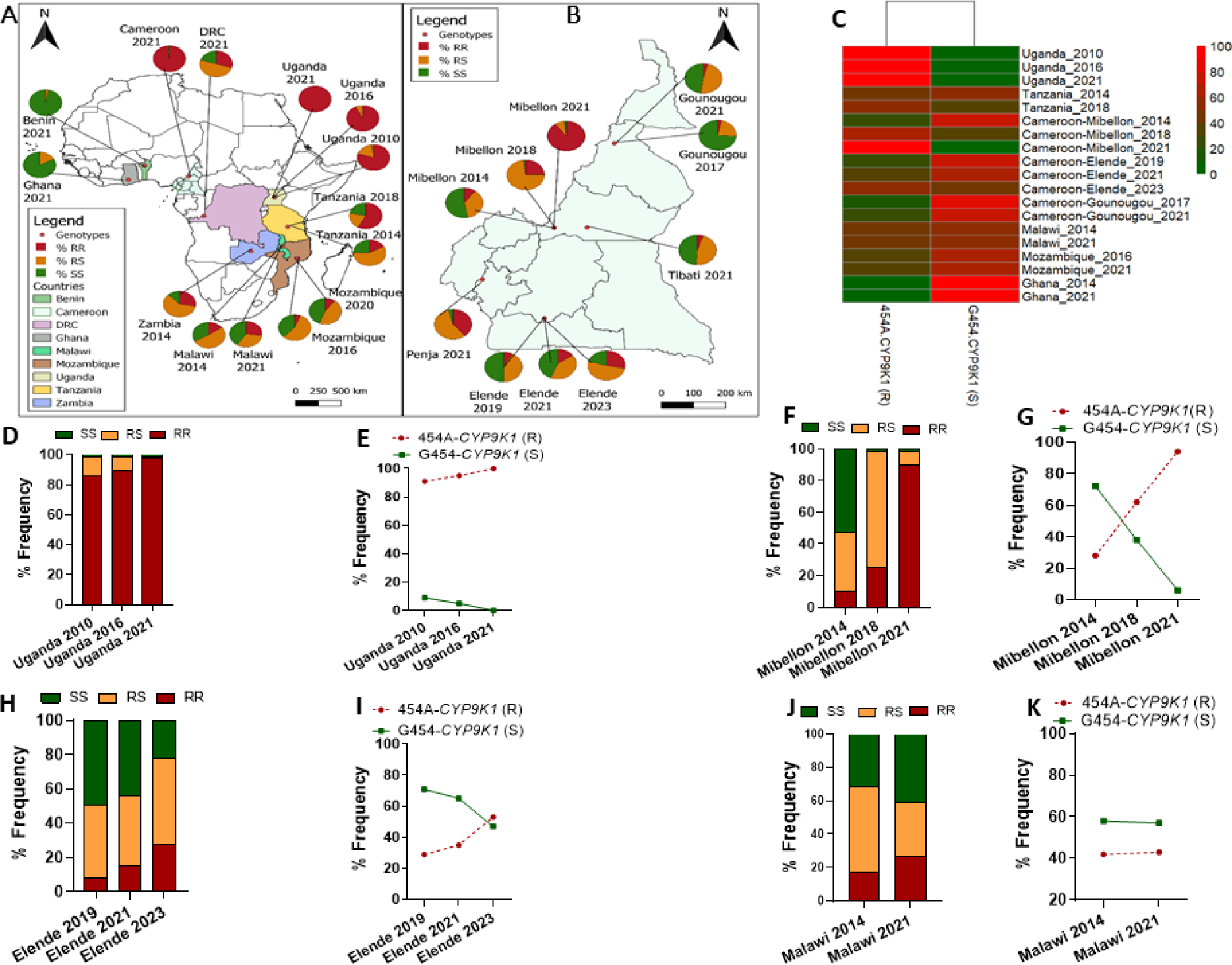
Spatio-temporal distribution of G454A-*CYP9K1* marker across Africa. (**A**) Africa-wide spatio-temporal percentage genotype distribution of G454A-*CYP9K1* marker, (**B**) Cameroon-wide spatio-temporal percentage genotypes distribution of *CYP9K1-G454A* marker, (**C**) Heat map showing Geographical and temporal evolution of *CYP9K1* percentage alleles frequency in *An. funestus* mosquito populations across Africa, showing increase in the frequency of the mutant 454A-*CYP9K1* allele in Uganda, Tanzania and Cameroon with time, presence but no change in frequencies of both alleles in Malawi and Mozambique with time and very low frequency of the mutant-type 454A-*CYP9K1* allele in Ghana (**D**) Temporal genotype distribution of G454A-*CYP9K1* marker in Uganda showing fixation of the mutant 454A/A-*CYP9K1* genotype in Uganda 2021, (**E**) Temporal evolution of *CYP9K1* alleles frequencies in Uganda showing fixation of mutant 454A-*CYP9K1* (R) allele and reduction to zero of the wild-type G454-*CYP9K1* (S) allele (**F**) and (**G**) are respectively same as (**D**) and (**E**) in Cameroon Mibellon showing increase in frequency of mutant 454A/A-*CYP9K1* genotype and 454A-*CYP9K1* allele with time and decrease in the frequency of the wild G454-*CYP9K1* allele with time. (**H**) and (**I**) are respectively same as (**D**) and (**E**) in Cameroon Elende showing increase in frequency of mutant 454A/A-*CYP9K1* genotype and 454A-*CYP9K1* allele with time and decrease in the frequency of the wild G454-*CYP9K1* allele with time. (**J**) and **(K)** are respectively same as (**D**) and (**E**) in Malawi showing increase in the frequencies of both the mutant 454A/A-*CYP9K1* and wild G/G454-*CYP9K1* genotypes with time (**J**) and no change in the frequencies of both the mutant-type 454A-*CYP9K1* and wild G454-*CYP9K1* alleles in Malawi between 2014 and 2021 (**K**).

Investigation of the distribution of this marker across Cameroon revealed its circulation across the country with varying frequencies according to regions (Fig 5B). Genotyping samples from Penja 2021 (Littoral region) revealed the marker was present at moderate frequency, 39% for the homozygote mutant genotype (RR) and very low frequency of 7% of the homozygote wild genotype (SS). The homozygote mutant genotype (RR) was detected at moderate frequency of 27% in Elende 2023 (Centre region). Assessing its distribution in other regions revealed the homozygote mutant genotype (RR) at very low frequencies 5% and 3% in Tibati 2021 (Adamawa region) and Gounougou 2021 (North region), respectively. The heterozygote (RS) and homozygote wild genotypes (SS) were the most abundant in Tibati 46% and 49%, respectively and in Gounougou 49% and 48%, respectively (Fig 5B).

Percentage allele frequency assessment revealed a high frequency of the mutant allele in samples from Eastern Africa (Uganda since 2010, Tanzania 2018) and Central Africa (Cameroon Mibellon 2021 and Elende 2023), moderate frequencies of both alleles in Southern Africa (Mozambique 2021 and Malawi 2021) and high frequency of wild allele in West Africa (Ghana 2021) (Fig 5C).

Temporal analysis of Ugandan samples (Eastern Africa) revealed increase in the frequency of the homozygote mutant genotype (RR) from 80% in 2010 to 93% in 2016 and 100% in 2021 (Fig 5D). Similar pattern was observed in Tanzania, with an increase in the frequency of mutant genotype (RR) from 17% in 2014 to 58% in 2018, respectively. Analysis of allele frequencies revealed an increase in the resistant 454A-*CYP9K1* (R) allele from 91% to 95% to 100% in samples from Uganda collected in 2010, 2016 and 2021 respectively (Fig 5E). Similarly, samples from Tanzania registered an increase in the mutant allele frequency (R) from 46% in female mosquito samples collected in 2014 to 68% in samples collected in 2018.

Monitoring of samples from Mibellon district (Adamawa region, Cameroon) registered a remarkable increase in percentage frequency of the homozygote mutant genotype (RR) from 10% in samples collected in 2014 to 25% in 2018 samples and 90% in 2021 female *An. funestus* mosquito samples (Fig 5F), leading to a massive increase in the percentage frequency of the mutant allele (R) in Mibellon from 28% in 2014 samples to 62% in 2018 samples and 94% in 2021 samples (Fig 5G).

In the same line, female *An. funestus* samples from Elende district (Centre region, Cameroon) registered a continuous increase in the percentage frequency of the homozygote mutant genotype (RR) from 8% in 2019 to 15% in 2021 and now 27% in 2023 (Fig 5H). This resulted in an increase in the percentage frequency of the mutant 454A-*CYP9K1* (R) allele in Elende from 28.5% in 2019 to 35% in 2021 and recently 53% in (Fig 5I). In the contrary, the wild G454-*CYP9K1* allele frequency gradually decreased from 50% to 44% in 2021 and 20% in 2023 (Fig 5I). Similar pattern was recorded Gounougou district (North region, Cameroon), with an increase in the percentage frequency of the mutant allele (R) from 13% in samples collected in 2017 to 24% in 2021.

Temporal monitoring of samples from Southern Africa from Malawi revealed an increase in percentage frequency for the homozygote mutant genotype (RR) from 15% in 2014 samples to 27% in 2021 samples (Fig 5J). Similar to Mozambique where we observed slight increase in percentage frequency of the homozygote mutant genotype (RR) from 6% in 2016 to 10% in 2020 and. However, at the allelic level no significant change in the percentage frequency of the mutant allele (R) was recorded in Malawi, from 42% in 2014 to 43% in 2021 (Fig 5K), similar to Mozambique where the percentage frequency of the mutant allele (R) slightly reduced from 34% in 2016 to 33% in 2021.

## Discussion

Dissecting the genetic bases of insecticide resistance is imperative for the identification of molecular markers required for the design molecular diagnostics, which is indispensable for the detection and monitoring of genes driving insecticide resistance. In this study, we unwound the role of the P450 G454A-*CYP9K1* marker in resistance to pyrethroid insecticide in *An. funestus* by *in vitro* comparative metabolism assays and *in vivo* comparative transgenic fly approach, highlighting the contribution of allelic variation (G454A) of *CYP9K1* gene as a key mechanism driving pyrethroid resistance. This facilitated the design of simple DNA-based molecular diagnostics around the G454A-*CYP9K1* marker. Applying this assay in the field to tract resistance revealed the mutant-type (454A*-CYP9K1*) allele is under strong directional selection in central and eastern Africa, strongly associated to pyrethroid resistance in these regions and reducing the efficacy of control tools (LLINs).

### The 454A-*CYP9K1* haplotype selected in Eastern Africa has spread to Central Africa

The low genetic polymorphism of *CYP9K1* originally recorded only in Uganda (Eastern Africa) in 2014 is now common to Cameroon (Central Africa). The fact that the same haplotype is predominant in both regions suggests that this 454A-*CYP9K1* mutant allele (R) has spread from East to West across the Equatorial zone of the continent. This similar to recent observations made for 4.3kb transposon-based pyrethroid resistant allele in *An. funestus* [33] suggesting that there is extensive gene flow between these regions from Uganda to Cameroon in line with genetic structure patterns reported in this species [32,34]. In previous whole genome association studies and targeted enrichment with deep sequencing, we reported reduced genetic diversity on the X-chromosome spanning the *CYP9K1* locus and the selection of the same haplotype in samples permethrin alive and dead from Uganda collected in 2014 whereas Cameroon samples were still very polymorphic [31,32]. Moreover, the high genetic polymorphism of the *CYP9K1* gene observed in Malawi (Southern Africa), and in laboratory mosquito strains FANG (Laboratory susceptible) highlights the little role played by the resistant allele of this gene in these regions in 2014. Previous studies have reported rapid selection of resistance with evidence of marked increase in the frequency of the *An coluzzii CYP9K1* haplotype in Bioko Island, Equatorial Guinea between 2011 and 2015 following reintroduction of pyrethroid-based control interventions [35], and in Selinkenyi, Mali before and after 2006 bed net scale up in this locality [36].

Furthermore, genetic variability analysis identified a single haplotype of this gene (454A) selected to fixation in mosquito samples from Uganda and Cameroon underlining the alarming rate at which this resistant allele could be rapidly spreading from Eastern to Central Africa. Polymorphism studies on other *An. funestus* cytochrome P450 monooxygenases *CYP6P9a/b*, *CYP6Z1*, *CYP6Z3* and the and the glutathione S-transferase gene epsilon 2 loci (*GSTe2*) genes highlighted selection of resistant alleles for each of these genes [28,29,37–39]. Conversely, genetic diversity studies of *CYP6M2* and *CYP6P4* in *An. coluzzii* and *An. gambiae s.s.* and some *An. funestus* cytochrome P450 genes, *CYP6Z1*, *CYP6Z3* and *CYP325A* have been shown to confer pyrethroid resistance but with no selection of dominant allele [27,28,40,41].

### The highly selected 454A-*CYP9K1* mutant-type allele is a better pyrethroid metabolizer

Comparative metabolism of recombinant *CYP9K1* alleles towards pyrethroid revealed higher metabolism for the mutant-type 454A*-CYP9K1* allele towards type II pyrethroid (deltamethrin) where metabolism of type I (permethrin) was lower. These findings support the evidence that allelic variation is greatly impacting the metabolic activity of the *CYP9K1* gene and increase in resistance driven by this allele. We had previously reported that 454A-*CYP9K1* allele coupled with *An. gambiae CPR* was mainly a type II (deltamethrin) metabolizer [31]. Similarly, *An. coluzzii CYP9K1* was reported to deplete deltamethrin at a similar rate [35]. Moreover, *An. arabiensis CYP6P4* gene was reported to metabolize permethrin (type I pyrethroid) with no metabolic activity towards deltamethrin (type II pyrethroid) [42] further supporting differential metabolism potential of certain genes between type I and II pyrethroids. Nevertheless, 454A-*CYP9K1* mutant-type allele could contribute to permethrin resistance by sequestration mechanism preventing the insecticides from reaching its target in significant amounts or by further detoxification of toxic metabolites produced from the action of other detoxification enzymes like the *CYP6P5* or *CYP325A* reported to be overexpressed in the same regions like the *CYP9K1* [16,26]. Several studies have laid evidence on allelic variation as a major mechanism driving differences in metabolic efficiencies for resistant genes like the *CYP6P9a/b, CYP325A* and *GSTe2* [27,28,38,43]. However, comparative metabolism of *An. gambiae* I236M-*CYP6P4* allelic variants reported both alleles were metabolisers of type I (permethrin) and II (deltamethrin) pyrethroids with no significant difference in the metabolic activity of both alleles towards pyrethroids [44].

### Allelic variation in *CYP9K1* combines with its over-transcription to confer higher pyrethroid resistance intensity

This study revealed that although both alleles (G454A) can confer resistance independently to both type I (permethrin) and type II (deltamethrin and alphacypermethrin) pyrethroids, greater resistance level was registered for flies expressing the mutant-type 454A-*CYP9K1* allele compared to those expressing the wild-type G454-*CYP9K1* allele. This enumerates the strong contribution of allelic variation (G454A) in addition to overexpression to confer even higher resistance levels in the flies carrying the mutant-type 454A-*CYP9K1* allele. Previous findings have reported transgenic *Drosophila* flies independently expressing the resistant alleles of the duplicated *CYP6P9a* and *CYP6P9b* genes to confer higher resistance level to pyrethroid,[28,29] and cross resistance to carbamate insecticide [43]. These findings correlates with studies of Riveron et al. (2014) [38] who reported that a single mutation L119F in *GSTe2* gene from *An. funestus* population is responsible for high resistance to both DDT and cross resistance pyrethroid [26,38]. Also, transgenic expression of brain-specific P450 *CYPBQ9* of *Tribolium castaneum* in *D. melanogaster* resulted in increased tolerance and lower mortality to deltamethrin insecticide compared to control flies [45]. Transgenic expression of other cytochrome P450s *CYP6G1*, *CYP6G2* and *CYP12D1* independently in *D. melanogaster* flies were shown to confer increased survival to at least one class of insecticide among DDT, nitenpyram dicyclanil and diazinon compared to control flies [46]. However, flies over-expressing *CYP6A2*, *CYP6A8*, *CYP6T3* or *CYP6A19* recorded no increased survival on any insecticide [46].

### Novel G454A-*CYP9K1* diagnostics are reliable DNA-based tools to detect and track the spread of metabolic resistance to pyrethroid insecticides

The design of this new cytochrome P450 DNA-based diagnostic tool G454A-*CYP9K1* AS-PCR and *CYP9K1*-LNA TaqMan assay greatly contributes to the efforts to detect and monitor the spread of metabolic resistance in the field. This marker correlates strongly with pyrethroid resistance using hybrid strains, as well as with field strains. This strong correlation with resistance to pyrethroid insecticide indicates allelic variation mechanism significantly contribute to the observed resistance driven by *CYP9K1*. This newly designed *CYP9K1* assay will supplement other known P450-based assays already in use, the *CYP6P9a* and *CYP6P9b* in *An. funestus* currently used to track resistance in Southern Africa [15,16]. Additionally, DNA-based diagnostic tools of metabolic resistance were previously designed for a 6.5kb structural variant insertion between *CYP6P9a/b* genes shown to contribute to resistance in Southern Africa [30], the recently detected 4.3kb transposon-based marker in East/Central African An. funestus populations [33] and the glutathione S-transferase gene epsilon 2 loci (*GSTe2*) in *An. funestus* conferring pyrethroid/DDT resistance and currently used to track resistance in West and Central Africa [38].

### Mutant-type 454A-*CYP9K1* allele reduces the efficacy of pyrethroid nets

The G454A molecular marker of *CYP9K1* significantly impacts the efficacy of some insecticide treated nets through cone test and experimental hut trials using free-flying F_0_ field *An. funestus* mosquitoes. PCR genotyping of the G454A-*CYP9K1* marker in alive and dead samples associates this reduced efficacy to the presence of 454A-*CYP9K1* resistant allele. This study carried out on field *An. funestus* (F_0_) samples in semi-field conditions spotlights the increasing challenge imposed by metabolic resistance on malaria vector control, contributing recent increase in malaria incidence and mortality in Africa [1]. Previous studies have also associated reduction of pyrethroid insecticide treated nets efficacy to metabolic resistant *An. funestus* markers *CYP6P9a*, the 6.5kb structural variant insertion and *CYP6P9b* [13,15,16,30]. Nevertheless, nets treated with the synergist PBO were shown to be much more effective than conventional pyrethroid-only nets in killing mosquitoes in populations where the G454A-*CYP9K1* mutant allele is present. Therefore, the use of PBO bed nets should be prioritized in the regions where this mutation is present.

### Africa-wide distribution pattern of G454A-*CYP9K1* supports the existence of barriers of gene flow in *An. funestus* across the continent

Assessment of spatio-temporal distribution of *CYP9K1*-G454A marker revealed increase in the frequency of the mutant-type allele with time in Eastern Africa (Tanzania and Uganda) and Central Africa (Cameroon). This highlights the intense selection of this allele in the field, mainly driven by the high selective pressure mostly from the scale up of insecticide-based interventions. This intense selection is more evident in Cameroon as the frequency of the 454A-*CYP9K1* moved from 28.5% in 2014 to 94% in 2021 highlighting the imperative needs to implement a robust surveillance of resistance in the field notably for new insecticides that are currently been introduced such as chlorfenapyr. The near fixation of 454A-*CYP9K1* in Cameroon (Mibellon), provides evidence of the major resistance role played by this gene in these localities. Transcriptomic and genomic studies have highlighted this gene as a potential candidate driver of resistance in these localities but requiring functional validation for further evidences [9,31,32]. Contrastingly, no evidence of directional selection of this mutant-type 454A-*CYP9K1* allele was observed in Southern Africa (Malawi and Mozambique) and Western Africa (Benin and Ghana), marked by the unchanging frequency of this mutant allele between 2014 and 2021 in these localities. This pinpoints the very little role played by this gene in these regions, with evidence that resistance in these regions is mainly driven by other genes notably the duplicated *CYP6P9a/b* [15,16,28,32,43]. The striking regional compartmentalisation of metabolic resistance markers in African populations of *An. funestus* highlights the challenge that barriers of gene flow may pose to future interventions such as gene drive or sterile insect techniques (SIT) which require these interventions to be implemented in different regions in parallel to overcome these barriers of gene flow.

### Conclusion

This study has demonstrated that allelic variation of *CYP9K1* gene is a major mechanism driving pyrethroid resistance in East and Central Africa. We showed that the resistant 454A-*CYP9K1* allele is a major driver of pyrethroid resistance illustrated by its close to fixation selection across these regions. The simple DNA-based molecular diagnostic designed here is a robust tool to detect and track the G454A-*CYP9K1* resistance in the field and assess it impact.

Interestingly, this resistant 454A-*CYP9K1* allele was shown to reduce efficacy of pyrethroid treated LLINs, but PBO-based nets were shown to have high efficacy even with the samples harbouring the mutant allele. This study offers a new DNA-based assay to track resistance in the field and improve resistance management strategies.

## Acknowledgements

We are thankful to the inhabitants of the collection sites for their support during the study. Special thanks to Helen Irving (LSTM) for reagents ordering and for handling the sequencing and to Dr. Mark Paine (LSTM) for the technical platform used for *in vitro* metabolism assays. The authors are also grateful to colleagues from CRID, Cameroon and vector biology department of Liverpool school of tropical medicine (LSTM), University of Liverpool for the support during the study.

## Data Accessibility

The *CYP9K1* sequences for 2014 and 2020 *An. funestus* samples generated during this study have been submitted to GenBank database (accession numbers: PP701029-PP701270).

## Author contributions

Design and conceptualization of the study: CSW. Methodology: CSDT, MFMK, MT, AM, LMJM, MJW, JH, SSI, and CSW. Sample collection: CSDT, MT, RFT, MG, and CSW. Investigation: CSDT, MFMK, AM, and NMTT. Visualization: CSDT, MT, JH, SSI, and CSW. Supervision: MT, LMJM AM, SSI, MHD, and CSW. Writing-Original draft: CSDT and CSW. Writing-review and editing: CSDT, MFMK, MT, AM, LMJM, JH, SSI, MHD, and CSW.

## Materials and Methods

### Mosquito samples collection and rearing

Mosquito samples used for this study were collected across different geographical regions in sub-Saharan Africa. Field mosquitoes *An. funestus* were collected from Eastern Africa Uganda, in Tororo district (0^◦^45ʹN, 34^◦^5ʹE) in March 2014 [52] and in Mayuge district (0^◦^23010.8ʹN, 33^◦^37016.5ʹE) in September 2020 [9], Southern Africa Malawi, Chikwawa district (12°19ʹS, 34°01ʹE) in January 2014 [53] and in June 2021 [11], Central Africa, Cameroon Mibellon (6°46′N, 11°70′E) in February 2015 [54] and in October 2020. and Elende districts (3°41′57.27’N, 11°33′28.46ʹE) in April 2019 [55], and in Western Africa, Ghana in Atatam village close to Obuasi district (06°17.377′ N, 001°27.545′ W) in July and in October 2021 [5]. The two *An. funestus* laboratory colonies used in this study were the FANG strain (fully susceptible to all insecticide classes) originated from Calueque district of Southern Angola (16°45ʹS, 15°7ʹE) and had been colonised in laboratory since 2002 [47]. and the FUMOZ (Pyrethroid resistant) colony derived from southern Mozambique that is highly resistant to pyrethroids and carbamates [47].

Adult Blood-fed female *An. funestus* were collected indoor in houses using electric aspirators between 06:00AM-10.00AM in each location. Females were left to become fully gravid within 4 days and then introduced into 1.5 ml Eppendorf tube to lay eggs following the forced-egg laying protocol [56]. The eggs were allowed to hatch and the larvae were reared to generate F_1_ adults used WHO bioassays. The field mosquito populations samples have been shown to be resistant to pyrethroids insecticide [5,8,9,11,54,55].

### Crossing between field and laboratory mosquitoes to generate hybrid strains

To segregate the various *CYP9K1* genotypes, two reciprocal crosses were generated in the insectary of the centre for research in infectious diseases (CRID). The first crossing was made between the female laboratory susceptible strain FANG and field male resistant strains *An. funestus* from Mibellon, Cameroon (FANG x Mibellon) and the second crossing was set up between the female laboratory strain FANG and the male resistant field strain from Mayuge, Uganda (FANG x Mayuge). These crossings were set up as described previously [51]. To achieve this, 100 F_1_ adult male *An. funestus* mosquitoes from Mayuge and from Mibellon were separately crossed with 120 females FANG. The crossings were followed up to F_3_ generation for FANG x Mibellon and the F_5_ generation for FANG x Mayuge. These crosses were used to make the correlation between *CYP9K1*-G454A mutation and resistance to pyrethroid.

### WHO insecticide susceptibility tests

The resistance profile to public health insecticides was determined using the WHO susceptibility bioassays protocol [57]. About 4 replicates each containing at least 20-25 female mosquitoes (either from field or hybrids from crossing) aged 2-5 days old were exposed to the insecticide(s) for either 10 minutes, 30 minutes or 1 hour, then transferred to holding tubes and provided with 10% sucrose solution. Four replicates of 20 mosquitoes each exposed to non-impregnated papers were used as control groups. The knockdown effect was recorded 1 hour after exposure time, while the mortality rates were recorded 24 hours post-exposure. These bioassays were conducted under standard conditions at 26 ± 2°C and 80 ± 10% relative humidity.

### Africa-wide temporal polymorphism analysis of *CYP9K1*

To assess the association between genetic diversity and resistance to insecticide, the polymorphism pattern of the *CYP9K1* gene was analysed across Africa firstly using target enrichment with deep sequencing (Sure Select) data for samples collected in 2014 across Africa and resistant to Permethrin insecticide. The sequence polymorphisms of a 1614 bp coding fragment spanning the full *CYP9K1* gene was analyzed between permethrin-resistant female *An. funestus* mosquitoes from each of the three countries (Uganda, Cameroon and Malawi) and in laboratory susceptible and resistant strains FANG and FUMOZ respectively. *CYP9K1* genetic diversity were retrieved from the SNP Multi-sample report file generated through Strand NGS 3.4 for each population. Bioedit [50] was used to input various polymorphisms in the vector base reference sequence using ambiguous letter to indicate heterozygote positions. Secondly, the full-length cDNA of this gene was amplified from permethrin-resistant *An. funestus* mosquitoes from Uganda 2020 (Mayuge), Cameroon 2020 (Mibellon), Ghana 2020 (Obuasi) and Malawi 2020 (Chikwawa). To provide additional contrast in the polymorphism analysis, full-length cDNA of *CYP9K1* was also amplified from laboratory fully susceptible strain FANG and the laboratory resistant strain FUMOZ. Full length coding sequences of *CYP9K1* was amplified separately from cDNA of 10 mosquitoes using the Phusion high fidelity DNA polymerase (Thermo Scientific) (primers sequences: Table S2). The PCR mixes comprised 3µl of 5x HF buffer (containing 1.5 mM MgCl_2_), 0.12µl of 25mM dNTPs, 0.51µl of 10mM forward and reverse primers, 0.15µl Phusion Taq, 9.71µl of deionised water and 1µl of DNA for a total 15µl reaction volume. Amplification was carried out using the following conditions: one cycle at 98°C for 1 min; 35 cycles each of 98°C for 20 s (denaturation), 60°C for 30 s (annealing), and extension at 72°C for 2 min; and one cycle at 72°C for 10 min (final elongation). PCR products were gel-purified using the QIAquick Gel Extraction Kit (Qiagen, Hilden, Germany), and ligated into pJET1.2/blunt cloning vector using the CloneJET PCR Cloning Kit (Fermentas). The recombinant *CYP9K1-pJET1.2* were used to transform cloned *E*. *coli DH5α* cells, recombinant plasmids miniprepped using the QIAprep Spin Miniprep Kit (Qiagen) and sequenced on both strands using the *pJET1.2* specific primers pJET1.2F and R primers provided in the cloning kit (Table S2). Sequences analysis involved manual examination using BioEdit version 7.2.3.0 [50] and aligned by multiple sequence alignments using ClustalW [58]. Population genetic parameters were assessed using DnaSP version 6.12.03 [49]. A haplotype network was built using the TCS program [59] to compare different haplotypes and a maximum likelihood phylogenetic tree was constructed using MEGA X [48].

### *In vitro* comparative assessment of metabolic activity of G454A-*CYP9K1* allelic variants Cloning and heterologous expression of recombinant *CYP9K1* alleles in *E*. *coli*

Recombinant enzymes of both alleles of *CYP9K1* were expressed as previously described for other P450s with slight modifications [27–29]. Expression plasmids *pB13::ompA+2-CYP9K1* for both alleles of *CYP9K1*: mutant-type (454A) allele from Uganda and Cameroon (454A-*CYP9K1-UGA*) and the wild-type (454G) allele from FANG (G454-*CYP9K1-FANG*) were constructed by fusing cDNA fragment from a bacterial ompA+2 leader sequence with its downstream ala-pro linker to the NH_2_-terminus of the P450 cDNA, in frame with the P450 initiation codon, as previously described [60], this was then cloned into *NdeI* and *EcoRI* linearised *pCW-ori+* expression vector [61]. To express the two recombinant alleles of *CYP9K1*, each allele was coupled with *An. funestus* cytochrome P450 reductase cloned in *pACYC* plasmid initially fused to the *peIB* leader sequence and used to co-transform *E. coli JM109* cells [62]. Membrane expression, preparations and measurement of P450 content were carried out as previously described [28,29,63] with slight modifications.

### Comparative assessment of pyrethroids metabolism of recombinant *CYP9K1* alleles

Metabolism assays using recombinant enzymes of *CYP9K1* allelic variants (G454 and 454A) with permethrin (type I pyrethroid insecticide) and deltamethrin (type II pyrethroid) were conducted following protocols described previously with some modifications [28,29,31]. To perform comparative metabolism assay, 0.2M KH2PO4 and NADPH-regeneration components (1mM glucose-6-phosphate, 0.25mM MgCl_2_, and 0.1mM NADP and 1U/ml glucose-6-phosphate dehydrogenase) were added to the bottom of 1.5ml tube chilled on ice. Membrane expressing recombinant G454-*CYP9K1* (wild-type) and 454A-*CYP9K1* (mutant-type) alleles, *AfCYPR*, reconstituted cytochrome b_5_ were added to the side of the tube and pre-incubated for 5 minutes at 30°C, with shaking at 1,200 rpm to activate the membrane. 20μM of test insecticide was then added into the final volume of 0.2ml (∼2.5% v/v acetonitrile), and reaction started by vortexing at 1,200 rpm and 30°C for 1 hour. Reactions were quenched with 0.1ml ice-cold acetonitrile and incubated for 5 more minutes. Tubes were then centrifuged at 16,000 rpm and 4°C for 15 minutes, and 150μl of supernatant transferred into HPLC vials for analysis. Reactions were carried out in triplicates with experimental samples (+NADPH) and negative controls (-NADPH). 100μl of sample was loaded onto an isocratic mobile phase (90:10 v/v acetonitrile to water) with a flow rate of 1ml/min, monitoring wavelength of 226nm and peaks separated with a 250mm C18 column (Acclaim 120, Dionex) on Agilent 1260 Infinity at 23°C. Enzyme activity was calculated as percentage depletion (the difference in the amount of insecticide remaining in the +NADPH tubes compared with the –NADPH) and a t-test used for statistical analysis.

### Comparative *in vivo* assessment of the ability of *CYP9K1* allelic variants to confer pyrethroid resistance using GAL4/UAS transgenic expression in *Drosophila*

Transgenic *D. melanogaster* flies expressing *CYP9K1* were used to investigate whether the over-expression of this gene could independently confer resistance to pyrethroid insecticide. Furthermore, to assess whether allelic variation in the *CYP9K1* significantly contributed to the resistance level, a mutant-type (Uganda) and a wild-type (FANG) alleles of this gene were independently expressed in *D. melanogaster*, using the GAL4/UAS system.

### Cloning and construction of transgenic lines

Two *CYP9K1 Drosophila melanogaster* transgenic lines, a mutant-type field line (*454A-CYP9K1-UAS-UGA*) and a wild-type laboratory line (*G454-CYP9K1-UAS-FANG*) were constructed. Cloning techniques, construction of transgenic flies and contact bioassays with transgenic flies was proceeded as described in previous studies [28,29,38]. The full-length *CYP9K1* was amplified from miniprep templates of the corresponding dominant alleles, using Phusion high fidelity DNA polyme rase (Thermo Scientific) and primers containing restriction enzyme sites for *EcoRI* and *XbaI* (Table S1). The PCR product was gel-purified using the QIAquick Gel Extraction Kit (Qiagen, Hilden, Germany), and ligated into pJET1.2/blunt cloning vector using the *CloneJET* PCR Cloning Kit (Fermentas). The *CYP9K1* alleles were then digested from the pJET1.2/blunt vector using restriction enzymes *EcoRI* and *XbaI*, purified and cloned in the *pUASattB* expression vector. These constructs were used to generate the transgenic lines 454A*-CYP9K1-UAS-UGA* and G454*-CYP9K1-UAS-FANG*. The primers used are listed in Table S2. Ubiquitous expression of *CYP9K1* candidate alleles in the transgenes in adult F1 progeny (the experimental group) was achieved after crossing homozygote males (UAS line with gene of interest) with virgin females from the driver strain Actin5C-GAL4. For control group, flies with the same genetic background as the experimental group but devoid of the UAS and gene of interest (pUASattb-*CYP9K1* insertion) construct were crossed with the driver Actin5C-GAL4 lines to generate null-Actin5C-GAL4 lines without the gene. All flies were maintained at 25°C in plastic vials with food.

### Insecticide contact Bioassays with transgenic flies

*Drosophila melanogaster* flies expressing the *CYP9K1* mutant-type (454A*-CYP9K1-UAS-UGA*) and wild-type (G454*-CYP9K1-UAS-FANG*) alleles were used for insecticide contact bioassays. Insecticide papers (2% permethrin, 0.2% alphacypermethrin and 0.2% deltamethrin-impregnated) filter papers were prepared in acetone and Dow Corning 556 Silicone Fluid (BDH/Merk, Germany) and kept at 4°C prior to bioassay as described previously [29]. These papers were rolled and introduced into 45cc plastic vials. The vials were then plugged with cotton wool soaked in 5% sucrose. 20-25 (2-4 days old post-eclosion *D. melanogaster* females) were selected for the bioassays and introduced into the vials. Mortality plus knockdown was scored after 1 hour, 2hours, 3hours, 6hours, 12hours and 24hours post-exposure to the discriminating doses of the insecticides. For each assay, at least six replicates were performed and t-test was used to carry out statistical analysis of mortality plus knockdown obtained between experimental groups and control.

### qRT-PCR validation of over-expression of transgenes

To confirm the expression of the candidate genes in the experimental flies and devoid in the control groups, qRT-PCR was carried out as described previously for other genes [29,64]. RNA was extracted separately from three pools of 5 F1 generation of experimental and control flies and used for cDNA synthesis. qRT-PCR for the *CYP9K1* was conducted using primers (primers listed in Table S3), with normalization using housekeeping genes *RPL11*.

### Design of DNA-based genotyping assay around the G454A marker of *CYP9K1*

To facilitate the detection and monitoring of resistance driven by the *CYP9K1* gene, two simple DNA-based PCR diagnostic assays; an allele specific PCR (AS-PCR) assay and a probe-based locked nucleic acid (LNA) assay were designed around the G/C-*CYP9K1* nucleotide SNP at position 1361 in the coding sequence of *CYP9K1* gene.

### Allele specific PCR (AS-PCR) design for G454A-*CYP9K1* mutation

The *CYP9K1* AS-PCR assay mainly reside on the use of two pairs of primers manually designed, a pair of outer primers made of an outer forward primer (9K1_OF: 5’-ACTGGACC GATGATGATTTGAC-3’) and an outer reverse primer (9K1_OR: 5’-ATCCAGAAGCC CTT CTCTGC-3’) and a pair of inner primers designed to match the mutation, comprised of an inner forward primer (9K1_IF: 5’-GGATCGTTTCTGG CCGGAACGGTTGG**C**-3**’**) and an inner reverse primer (9K1_IR: 5’-TATCGATCGGT GTCGGGCTGTC CGCTC-3’) as previously described [65]. An additional mismatched nucleotide (underlined) was added in the third nucleotide from the 3′ end of each inner primer (9K1_IF: 5’-GGATCGTTTCTGGCCGG AAGGTT<u>G</u>G**C-3’** and 9K1_IR: 5’-TATCGATCGGT GTCG GGCTGTCCG <u>C</u>T**C-3’** to enhance the specificity.

### Locked nucleic acid (LNA) assay design for genotyping of G454A-*CYP9K1* marker

A locked nucleic acid was also designed around the G454A*-CYP9K1* marker to provide alternative and additional PCR-based diagnostic assay for *CYP9K1* to track resistance in the field. This assay relies mainly on a pair of LNA probes and primers designed around this G454A-*CYP9K1* mutation. LNA primers for this assay were designed to amplify 95 bp product surrounding the G454A-*CYP9K1* codon, and LNA probes were designed according to previously suggested parameters with slight modifications [66]. The *CYP9K1*-LNA probes (LNA9K1-Gly: Hex: TCCGG+T+C+CGAAC and LNA9K1-Ala: Fam: TCCGG+ T+G+CG+AA) and primers (LNA-9K1F: 5’-CGTGATCCGCAACTGTTTC-3’ and LNA-9K1R: 5’-GTAAGGATGGACGCGGTATC-3’) were synthesised by integrated DNA technologies (IDT) (http://biophysics.idtdna.com/) (Table S4). Each assay was prepared to contain a final concentration of 1× PrimeTime Master Mix (IDT) or 1× Luna Universal qPCR Master Mix (NEB), 0.1 μM for each of the two probes (LNA9K1-Gly: Hex and LNA9K1-Ala: Fam), 0.2 μM of primers (LNA-9K1F and LNA-9K1R) in a total reaction volume of 10 μl with 1 μl of DNA template. Reactions were set up in optical PCR tubes and run on an AriaMX Real-Time qPCR cycler (Agilent, USA) with Fam and Hex filters. The G454A*-CYP9K1*-LNA assay consisted of a 3 minute denature at 95°C; 20 cycles of denaturation for 15 seconds at 95°C, annealing for 30 seconds at 66°C; 23 cycles of denaturation for 10 seconds at 95°C, annealing for 20 seconds at 58°C, and an extension of 10 seconds at 72°C.

### Correlation between the G454A-*CYP9K1* mutation and resistance phenotype

Two hybrid strains (FANG x Mibellon and FANG x Mayuge) and field strains (Elende 2022) were used to assess genotype phenotype association. This mutation was genotyped in alive and dead samples generated from FANG x Mayuge at F_5_ generation bioassayed to permethrin and FANG x Mibellon F_3_ generation bioassayed to alphacypermethrin by WHO tube test. Genomic DNA was extracted following Livak protocol [67] and used for PCR genotyping using the newly designed assays. PCR genotyping was carried out using 10 mM of each primer and 1ul of genomic DNA as template in 15 μl reaction mix containing 10X Kapa Taq buffer A, 0.2 mM dNTPs, 1.5 mM MgCl2, 1U Kapa Taq (Kapa biosystems). The cycle parameters were: 1 cycle at 95 °C for 2 min; 30 cycles of 94 °C for 30 s, 59 °C for 30 s, 72 °C for 1 min and then a final extension step at 72 °C for 10 min. PCR products were separated on 1.5% agarose gel electrophoresis, stained with Midori Green advance DNA Stain (Nippon Genetics Europe GmbH) and visualised on a UV transilluminator. To make the correlation between the *CYP9K1* G454A marker and resistance in the field conditions, Elende (Cameroon) field mosquito samples (F_1_) generations collected in 2022 were used for bioassays with 0.05% alphacypermethrin insecticide. Genomic DNA was extracted from mosquito samples alive and dead after bioassays and used for *CYP9K1* genotyping using the newly designed *CYP9K1* AS-PCR assay. The various genotypes from alive and dead samples were used to make the correlation between this marker and resistance in the field mosquito samples.

### Assessment of the impact of G454A-*CYP9K1* marker on the efficacy of LLINs using Cone bioassays

The efficacy of some standard pyrethroid and PBO-pyrethroid was evaluated using F_1_ Elende field samples and following the WHO cone assay protocol [68]. The nets tested included; PermaNet 2.0 (Deltamethrin), Duranet (Alphacypermethrin), Royal sentry (Alphacypermethrin), Olyset (Permethrin), Olyset plus (Permethrin with PBO), PermaNet 3.0 top (Deltamethrin with PBO), PermaNet 3.0 side (containing only deltamethrin). Five replicates of 10 mosquitoes each were exposed to 30 cm x 30 cm pieces of each of the nets through the cone for 3 minutes and transferred into paper cups. The knockdown effect was recorded 1 hour after exposure and the mortality rate was recorded 24 hrs post-exposure. Controls comprise 4 replicates of 10 mosquitoes exposed to 2 pieces of untreated nets.

### Assessment of the impact of G454A-*CYP9K1* marker on LLINs through EHTs

The experimental hut study was performed in Elende (3°4157.27′N, 11°3328.46′E), a district in the Centre region of Cameroon, where 12 experimental hut stations were recently constructed with concrete bricks and a corrugated aluminium roof, following the specific designs elaborated for West African region experimental huts constructions [68]. Three treatments were compared in this study, comprised of an untreated polyethylene net; a standard net, Royal sentry (alphacypermethrin impregnated polyethylene net); and a PBO-net, PermaNet 3.0 (PBO + Deltamethrin incorporated into polyethylene net). To reflect a worn net, six 4 cm × 4 cm holes were made on each net according to WHO guidelines. Three adult volunteers were recruited from Elende village to sleep under the nets and collect mosquitoes in the morning. Each volunteer was provided a written consent to participate in this study and were also given chemoprophylaxis during the trial. Ethical approval for this study was obtained from the national ethics committee for health research of Cameroon (ID: 2021/07/1372/ CE/CNERSH/SP). Early in the morning, mosquitoes were collected using glass tubes from the room (the floor, walls, and roof of the hut), inside the bed net, and from the exit traps on the veranda. Surviving mosquitoes were provided with sugar solution and held for 24 hours in paper cups after which delayed mortality was assessed. Samples were recorded in observation sheets as dead/blood-fed, alive/blood-fed, dead/unfed, and alive/unfed. The effect of each treatment was expressed relative to the control by assessing the killing effect.

### Genotyping of G454A-*CYP9K1* marker in EHT samples

To assess the impact of the *CYP9K1-*mediated resistance to pyrethroids on the effectiveness of the insecticide-treated nets (Royal sentry, PermaNet 3.0), the newly designed AS-PCR assay was used to genotype a subset of each treatment mainly the dead and alive mosquitoes on the veranda, in the net and in the room.

### Africa-wide spatio-temporal assessment of the Spread of G454A-*CYP9K1* marker

To assess the spread and temporal changes in the frequency of the G454A-*CYP9K1* marker in Africa, *An. funestus* samples previously collected at different time points in the same localities across Africa [5,7,9,11,32,54,55], involving eastern Africa (Uganda 2010, 2016 and 2021 and Tanzania 2014 and 2018), central Africa (Cameroon (Mibellon 2016, 2018 and 2021, Elende 2019, 2021 and 2023, Gounougou 2017 and 2021, Tibati 2021, Penja 2021) and DRC 2021), southern Africa (Zambia 2014, Malawi 2014 and 2021 and Mozambique 2016 and 2020) and western Africa (Benin 2021 and Ghana 20121) were used for the study. Genomic DNA extracted from these samples were genotyped using the newly designed assay.

### Data Analysis

Statistical analysis was performed with Prism 8 (GraphPad Software, San Diego, California USA, www.graphpad.com) and alpha values for significance were taken at *P* < 0.05, with all confidence intervals (CI) at 95%. Student’s *t* test was used to compare two columns of data generated from metabolism assays and contact bioassays with transgenic *D. melanogaster* flies. Fisher’s exact test was used to determine whether any difference in proportion observed for the genotype contingency tables was significant. Statistical significance was denoted by asterisks: *P* > 0.05, \**P* < 0.05, \*\**P* < 0.01, and \*\*\**P* < 0.001. QGIS version 3.14.16 (www.qgis.org), was used to generate the map presenting the percentage frequency of various genotypes of G454A-*CYP9K1* marker per genotyping sites across Africa.

## Supporting Information

**S1 Fig. Schematic representation of polymorphism positions of *An. funestus CYP9K1* sequences from 2014 samples showing** (**A**) nucleotide changes, the guanine to cytosine nucleotide at position 1361 (highlighted in orange colour). (**B**) Polymorphic positions of *CYP9K1* amino acid sequences 2014 samples the G454A (in green).

**S2 Fig. Optimization of DNA-based diagnostic assay for G454A-*CYP9K1* genotyping (A)** Agarose gel image of G454A-*CYP9K1* AS-PCR showing homozygote mutant genotypes 454A/A-*CYP9K1* (RR, with band size 216 bp and a common band 639 bp) for all Uganda (Mayuge F_0_ 2021), homozygote wild genotype G/G454-*CYP9K1* (SS, with band size 434 bp plus a common band 639 bp) for FANG samples, **(B)** Dual color scatter plot of *CYP9K1* Probe-based locked nucleic acid (LNA) showing homozygote mutant genotypes 454A/A-*CYP9K1* (RR) clustered on the y-axis for all Uganda (Mayuge F_0_ 2021) samples, homozygote wild-type G/G454-*CYP9K1* (SS) genotypes clustered on the x-axis for all FANG samples, molecular weight Marker (MW), negative control (NC) and non-amplification (NA).

**Table S1. Summary statistics of the correlation between G454A-*CYP9K1* mutation and pyrethroid resistance phenotype.**

**Table S2. Summary statistics of the correlation between G454A-*CYP9K1* marker and efficacy of LLINs by WHO cone test and EHTs.**

**Table S3. Primers used for *CYP9K1* cDNA amplification and functional validation Table S4. Primers and probes used for G454A-*CYP9K1* marker genotyping**

